# Neuroimmune circuits involved in β-lactoglobulin-induced food allergy

**DOI:** 10.1101/2020.08.17.248310

**Authors:** Luísa Lemos, Helder Carvalho Assis, Juliana Lima Alves, Daniela Silva Reis, Maria Cecilia Campos Canesso, Thais Garcias Moreira, Barbara Kaori Sato Miranda, Luara Augusta Batista, Julia Ariana de Souza Gomes Lenzi, Muiara Aparecida Moraes, Luciana Melo Pereira, Daniele Cristina Aguiar, Bruno Rezende Souza, Denise Carmona Cara, Ana Cristina Gomes-Santos, Ana Maria Caetano Faria

## Abstract

Cow’s milk allergy is the most prevalent food allergy that usually begins early in life and β- lactoglobulin (BLG) is the milk component with the highest allergenicity. It has been described that ovalbumin (OVA)-induced food allergy in mice is associated with anxiety and aversive behavior. However, it is yet to be determined whether altered behavior is a general component of food allergy or whether it is specific for some types of allergens. Thus, we investigated behavioral and neuroimmune circuits triggered by allergic sensitization to BLG. We found a neuroimmune conflict between aversion and reward in a model of food allergy induced to BLG. Mice sensitized to BLG did not present aversive behavior when the allergen was used for sensitization and oral challenge. Mice allergic to BLG preferred to drink the allergen-containing solution over water even though they presented high levels of specific IgE, inflammatory cells in the intestinal mucosa and significant weight loss. When sensitized to OVA and orally challenged with the same antigen, mice had display neuron activation in the amygdala suggesting an anxiety-related sensation. On the other hand, OVA-sensitized mice showed preference to consume a mixture of BLG and OVA during oral challenge in spite of their aversion to OVA. Consumption of OVA-BLG solution was associated with neuron activation in the nucleus accumbens, suggesting a reward sensation. Thus, the aversive behavior observed in food allergy to OVA does not apply to all antigens and some allergens may induce preference rather than aversion. Our study provides new insights into the neuroimmune conflicts regarding preference and avoidance to a common antigen associated with food allergy.

## INTRODUCTION

Food allergy is an abnormal immune response against dietary antigens (Johansson et al., 2004). Among all food allergies, cow’s milk allergy is one of the most prevalent and β- lactoglobulin (BLG) is the primary allergen involved (Jo et al., 2014). Allergen-specific immunoglobulin (Ig) E produced by plasma cells in circulation binds to Fc∊RI receptors expressed on the membrane of mast cells and basophils, which upon a second allergen exposition, undergo a massive degranulation with the release of various inflammatory mediators that trigger the development of allergic reactions sometimes as severe as anaphylactic shock (Bischoff and Crowe, 2005). Importantly, a response triggered by the immune system during food allergy can affect the central nervous system through direct signals from sensory nerves present in the intestine (van der Kleij et al., 2010). Moreover, mast cells may establish direct contact with nerves through cell adhesion molecules type 1 (CADM-1) (Furuno et al., 2012; Hagiyama et al., 2011) and act directly on neurons of the enteric nervous system (Voisin et al., 2017a). As a response, neurons secrete mediators such as neuropeptides and neurotransmitters that act on cognate receptors expressed on immune cells involved in food allergy (Dantzer, 2018). Probably this bidirectional neuroimmunological interaction occurs early in the reaction having a great impact on allergic inflammation (Voisin et al., 2017b). Due to this complex net of neuroimmune signaling, symptoms of allergy may range from a slight inconvenience to life-threatening reactions such as anaphylactic syndrome associated with high degree of anxiety (Sicherer, 2003).

Anxiety sensation have been the most common behavioral pattern associated with food allergy (Costa-Pinto and Basso, 2012). Allergic patients in a state of anxiety often have their daily tasks compromised due to a feeling of eminent risk (Sampath et al., 2018). Our group showed that mice previously sensitized to ovalbumin (OVA) avoid drinking the allergen-containing solution even when it contains a palatable component such as saccharin (Cara et al, 1994). OVA-sensitized mice orally challenged with the allergen present increased levels of anxiety evidenced by shorter time of exploration in the open arms of an elevated plus maze (EPM) and strong activation of specific brain areas involved in anxiety such as the paraventricular nucleus of the hypothalamus (PVN) and the central nucleus of amygdala (CeA) (Salgado et al., 2004). Furthermore, it has been shown that sensitized mice avoid a compartment previously associated with presentation of the allergen to which they were sensitized (Costa-Pinto et al., 2005). In a model of food allergy to OVA, the expression of mRNA for the calcitonin gene-relate peptide (CGRP) was shown to be increased in the colon of mice whereas the distribution of nerve fibers did not change, suggesting that CGRP release may be increased during allergies (Lee et al., 2013). Furthermore, Basso and coworkers have shown that a Th2-associated response induces changes in central nervous system activity and behavior through an IgE-dependent mechanism (Basso et al., 2003), suggesting that IgE may play an important role in the aversive behavior observed in experimental food allergy. Consistent with this, passive transfer of hyperimmune serum or adoptive transfer of splenocytes from mice allergic to OVA into naïve mice transferred food aversion (Cara et al., 1997). However, little is known about the behavioral aspects of food allergy induced by other allergens. Thus, in the present study, we aimed to investigate whether mice with BLG-induced food allergy had altered behavior and the neuroimmune circuits underlying these putative behavioral changes. We found that mice sensitized to BLG developed a partial aversion to consumption of a BLG-containing solution that was further lost despite the high levels of anti-BLG IgE and the great weight loss. This is a remarkable distinct behavior when compared to OVA-sensitized mice challenged with OVA. Moreover, mice sensitized to OVA that received a mixture of BLG and OVA during oral challenge, preferred to drink this mixture, despite their aversion to OVA. This preference behavior was associated with neuron activation in the nucleus accumbens, suggesting that a reward sensation was triggered by consuming the allergen. Thus, different allergens induce different responses, which is likely a result from the cerebral region they activate.

## MATERIAL AND METHODS

### Animals

Male BALB/c mice at 7 to 8 weeks of age were obtained from Universidade Federal de Minas Gerais (UFMG, Brazil) animal facility and maintained under specific pathogen free condition. All procedures were in accordance with the ethical principles in animal experimentation, adopted by the Ethics Committee in Animal Experimentation of our institution (CEUA - UFMG) - Protocol 144/2019. Mice were kept in a temperature-controlled room with free access to water and standard chow diet up to the oral antigen challenge period (24 h prior to experiments).

### Mice sensitization and oral challenge

Food allergy was induced as previously described by Gomes-Santos (2015). Briefly, mice received intraperitoneally (i.p.) 0.2 ml saline (0.9%) solution containing 1 mg Al (OH)_3_ as an adjuvant and 20 µg BLG (Sigma, St. Louis, MO, USA). Control mice received 0.2 ml saline (0.9% p/v) solution containing 1 mg Al (OH)_3_. After 14 days, all mice were i.p. immunized with saline containing 20 μg soluble BLG. Control mice received only i.p. saline (0.9% p/v). Seven days later, all mice were orally challenged with 20% whey protein solution in the drinking bottle as their only source of liquid for a period of either 7 or 14 days. Solutions were replaced daily. Whey protein hydrolysate was obtained from EDETEC and contained 80% BLG. Alternatively, mice received either 1 mg/mL BLG, OVA (Sigma, St. Louis, MO, USA) or a mixture of both.

### Liquid intake

Liquid consumption was measured during experimental analysis by checking the remaining quantity of liquid in the bottle and the quantity offered in the previous day. The result obtained was the average consumption per mouse/day.

### Food-choice in a food-preference test

A food preference test was performed to evaluate the choice for BLG solution over water. Three days before the test was conducted, mice were individually separated in conventional cages from the animal facility for acclimation. On the same day, two identical, transparent, glass-tipped bottles containing water were placed for each animal to choose between the bottles. In the food preference day test, two bottles were placed per cage: one containing the antigenic solution (OVA or BLG or the mixture of both) and the other containing only water. Liquid consumption was measured every 4 hours during a 24-hour period, which occurred on the first day of the oral challenge. After each measurement, bottles were filled with the liquid, however the position was changed to avoid conditioning by their location in the cage.

### Measurement of serum specific anti-BLG antibodies

To measure the concentration of anti-BLG IgE, capture-enzyme-linked immunosorbent assay (ELISA) was applied, in which plates coated with rat anti-mouse IgE was used, as previously described by our group (Gomes-Santos et al., 2015). Briefly, serum samples were obtained from all mice groups after the last oral antigen exposure. Fifty µl total serum, biotinylated BLG, and HRP-labeled streptavidin were added individually to a 96-well plate. Subsequently, the reaction was developed by adding H_2_O_2_ with orthophenylenediamine (OPD, Sigma, St. Louis, MO, USA) as previously described (Russo et al., 2001). Results obtained were reported in arbitrary units using a positive reference serum (1000 U).

To measure the concentration of anti-BLG IgG1, plates were incubated with 100μl/well of a BLG solution (2μg/well) diluted in carbonate buffer pH 9.6 and kept overnight at 4° C. Next day, after three washes with PBS solution, plates were blocked by adding 200μl/well of a 0.1M phosphate buffer casein solution (pH 7.4) for at least 1 hour at room temperature. Diluted serum was added at 1: 400, and serial dilutions were performed. After plates were incubated at 37° C for one hour and washed, peroxidase-labeled anti-isotype antibodies (Goat anti-mouse IgG1-HRP; Southern Biotechnology Associates Inc.) were added at 1:15.000 dilution. The reaction was developed by adding H_2_O_2_ with OPD.

To measure the concentration of anti-BLG secretory IgA, small intestine from mice were collected and rinsed with 10mL cold 0.1M phosphate buffer solution (PBS, pH 7.4). Intestinal lavages obtained were centrifuged at 12,000g for 20 min at 4ºC. Supernatants were then collected and secretory IgA concentration was determined by ELISA (Gomes-santos et al., 2012).

### Measurement of proximal jejunum cytokines

Proximal jejunum samples were weighed and homogenized in PBS containing 0.05% Tween-20, 0.1 mM phenylmethylsulphonyl fluoride, 0.1mM benzethonium chloride, 10mM EDTA and 20 KIU Aprotinin A using a tissue homogenizer (100mg tissue/ml buffer). Suspensions were centrifuged at 12,000g for 20 min at 4ºC and the supernatants were transferred to microtubes and stored at −80ºC until analysis. Concentrations of IL-4, IL-5, IL-10 were measured by ELISA.

### Intraepithelial lymphocytes (IEL) counting

The intestinal epithelium was examined on histological slides stained with hematoxylin-eosin (H&E) in 20X magnification under an optical microscope. The IELs were identified by their characteristic localization: basal to the nuclei of the enterocytes and small clear halo of cytoplasm around their dense and regular spherical nucleus. For each fragment, 500 epithelial cells were counted, not including goblet cells, as described previously (Ferguson and Murray, 1971). The final number of IELs was expressed as IEL per 100 counted epithelial cells.

### Elevated Zero Maze (EZM)

The maze is a device made of acrylic plastic and has two open arms and two closed arms (Cruz et al., 1994) and elevated 100cm from the floor. The arms (open or closed) are positioned on opposite sides. In the EZM test, each mouse is placed individually in the central area of the maze, where they can explore its arms freely for 15 minutes. The time spent exploring the open and closed arms is recorded by video, as well as the number of entries and exits on each arm. The percentage of time spent in the open arms of the maze is inversely proportional to the level of anxiety in the test.

### Quantification of c-Fos in the brain by immunohistochemistry

Ninety minutes after exposure to the maze, mice were anesthetized with urethane and perfused transcardially with 4% paraformaldehyde in 0.1 M phosphate buffer (PB, pH 7.4). Brains were removed and fixed for two hours in paraformaldehyde and stored for at least 30 hours in 30% sucrose for cryopreservation. 40 μm-thick coronal sections were cut with a freezing microtome and were cryopreserved in duplicate in the cryostat. Sections were first processed for c-Fos labeling by immunohistochemistry, as previously described (Beijamini and Guimarães, 2006; de Oliveira et al., 2000). Finally, after dehydration in xylene diaphanization, slides were mounted in Entellan^®^.

### Histological analysis of proximal jejunum

Animals were sacrificed and their proximal jejunum collected and opened longitudinally for histological analyses. Proximal jejunum was fixed with 10% formalin in neutral buffer and embedded in paraffin. Histological sections were previously deparaffinized and then stained with H&E or Periodic acid-Schiff and analyzed by light microscopy (Olympus BX41). The histological changes were evaluated in a double-blinded fashion.

### Statistics

Results were reported as the mean ± standard deviation. One-way ANOVA with Tukey post-hoc analyses were used for multiple comparisons. P-values under 0.05 were considered significant as compared to the control group. Graphs and statistical analyzes were performed using GraphPad Prism version 6.00 for Windows (GraphPad Software, San Diego, CA, USA).

## RESULTS

### BLG induces food allergy

To investigate whether BLG induced food allergy, mice were first sensitized to BLG and then challenged with whey protein solution (containing 80% of BLG) for 7 or 14 days (Fig. 1A). We found that mice sensitized and challenged with whey protein at both experimental times lost weight (Fig. 1B), which is an important feature of food allergy. Similarly, a marked reduction of epididymal adipose tissue mass in mice challenged with whey protein was observed, whereas non-sensitized mice did not lose weight (Fig. 1C). We also quantified chow consumption to verify whether the weight loss related to a reduced consumption of chow and found that all groups had a similar chow consumption (Fig. 1D).

**Figure 1.**
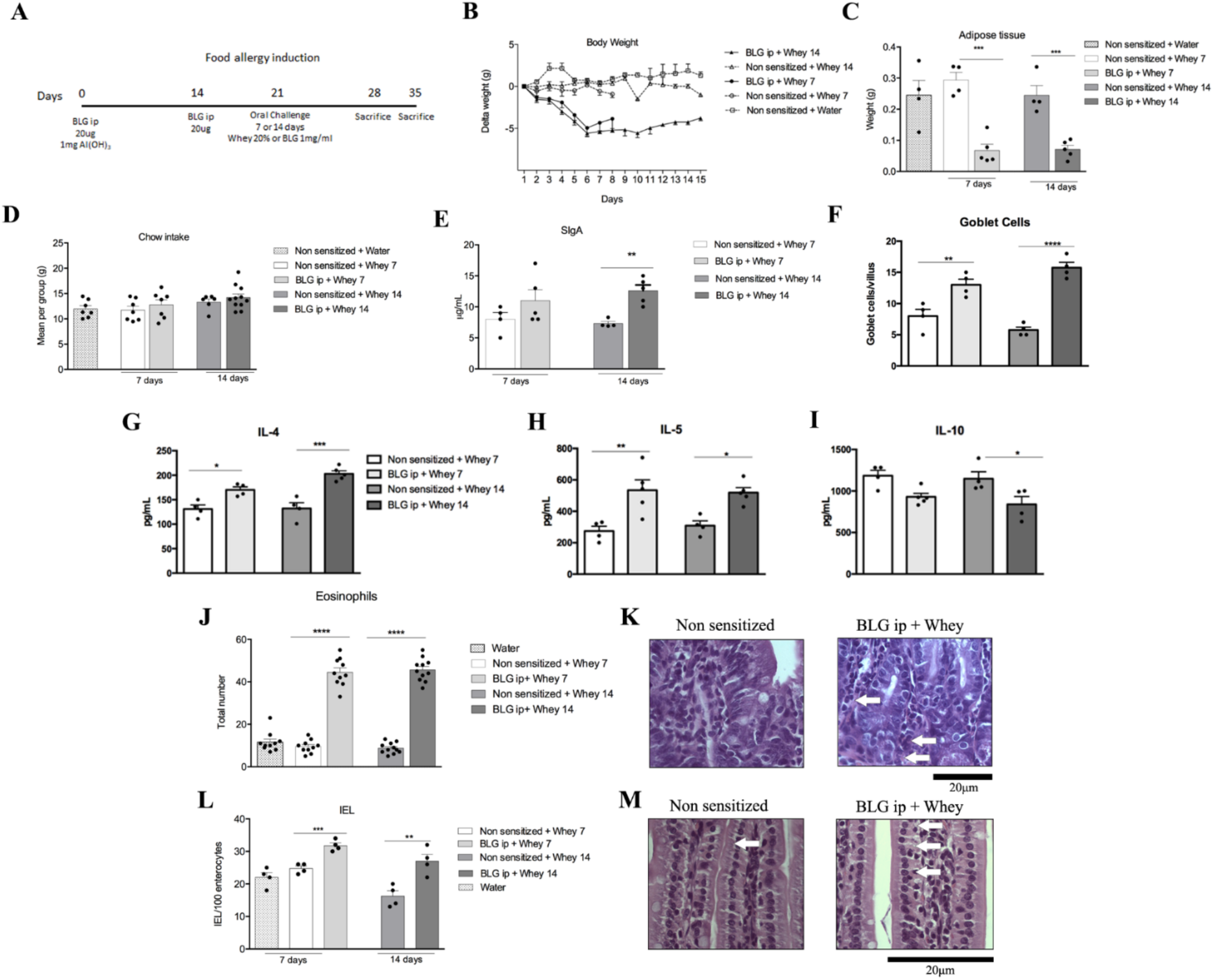
Food allergy induced to beta-lactoglobulin. A) Mice were sensitized with 20ug of BLG ip adsorbed in 1mg of Al (OH)3. After fourteen days, mice received a booster with BLG ip. One week later, mice were orally challenged with 20% Whey diluted in either water or BLG 1mg/ml for either 7 or 14 days being the only source of liquid, then mice were sacrificed. B) Body weight of mice represented by delta weight during 14 days of oral challenge. C) Weight of epididymal adipose tissue after oral challenge. D) Chow intake among groups during oral challenge. Data represent 2 independent experiments with 4 to 6 mice/group. Data represent the mean +- SEM. \*\*\**p*< 0.001. E) SIgA levels were measured in small intestine lavage by ELISA as previously described in methods. Data was represented as means ± SEM of non sensitized and sensitized mice in both 7 and 14 days of oral challenge with whey protein solution. F) Goblet cells of proximal jejunum were counted, and total number was expressed as mean ± SEM. ^*^*p* < 0.05, ANOVA-Tukey. Type-2 cytokines IL-4 (G), IL-5 (H) and IL-10 (I) were measured in proximal jejunum. J) Eosinophils counted in random fields of histology slides. K) Representative photomicrographs of H&E-stained eosinophils. L) Number of IEL counted in the proximal jejunum of mice after oral challenge in histology slides. M) Representative photomicrographs of H&E-stained IEL.

Next, we measured the levels of anti IgE specific in serum and small intestine secretory IgA (sIgA). We found that serum anti-BLG IgE were elevated in mice that were previously sensitized to BLG and subsequently challenged with whey protein solution, but not in non-sensitized mice (Fig. 1E). BLG-sensitized mice that received water throughout the experiment, but were not challenged orally with whey protein, had higher levels of anti-BLG IgE than non-sensitized animals, however, these levels were still significantly lower as compared to mice that were sensitized to BLG and orally challenged with whey protein (Fig. 1E). Moreover, mice sensitized to BLG and orally challenged with whey protein for 14 days had higher levels of anti-BLG IgE than mice challenged for 7 days (Fig 1E). Furthermore, sIgA in small intestinal lavage fluids was increased in allergic mice when they were challenged for 14 days, but not for 7 days, with whey protein (Fig. 1F).

We also measured cytokines known to participate in allergic responses such as IL-4, IL-5 and IL-10 in the proximal jejunum of mice. Oral challenge with whey protein in sensitized and orally challenged mice induced increased levels of IL-4 (Fig. 1G) and IL-5 (Fig. 1H) during oral challenge, but IL-10 levels were decreased only after 14 days of oral challenge with whey protein (Fig. 1I). Moreover, ingestion of whey protein induced an increase in goblet cell numbers (Fig. 1J), which produce and secrete mucus (Birchenough et al., 2016), and inflammatory cell infiltration in the gut of sensitized mice 7 and 14 days after oral challenge. As part of this inflammatory reaction, the number of eosinophils was higher in sensitized mice challenged with whey protein (Fig. 1K, L). Intraepithelial lymphocytes (IEL) were also increased in the intestinal mucosa of BLG-sensitized mice that were challenged orally for 7 and 14 days (Fig. 1M). Thus, mice sensitized with BLG and orally challenged with whey protein develop food allergy.

### Mice allergic to BLG develop a partial aversive behavior

The aversive behavior occurs when mice avoid consuming solutions or diets containing the antigen to which they are allergic (Cara et al., 1994). To investigate whether mice allergic to BLG showed differences in consumption of a solution containing whey protein, which a reduction in intake would indicate aversion, we assessed their daily consumption of whey protein and found that both non-allergic (non-sensitized) and allergic (BLG-sensitized) mice consumed significantly more whey protein solution than the control group that was only exposed to water (Fig. 2A). However, allergic mice consumed less whey protein solution than non-allergic mice during 12 days, suggesting that mice allergic to BLG developed an aversive behavior. Moreover, this aversive behavior was lost after 12 days of challenge (Fig. 2A).

**Figure 2.**
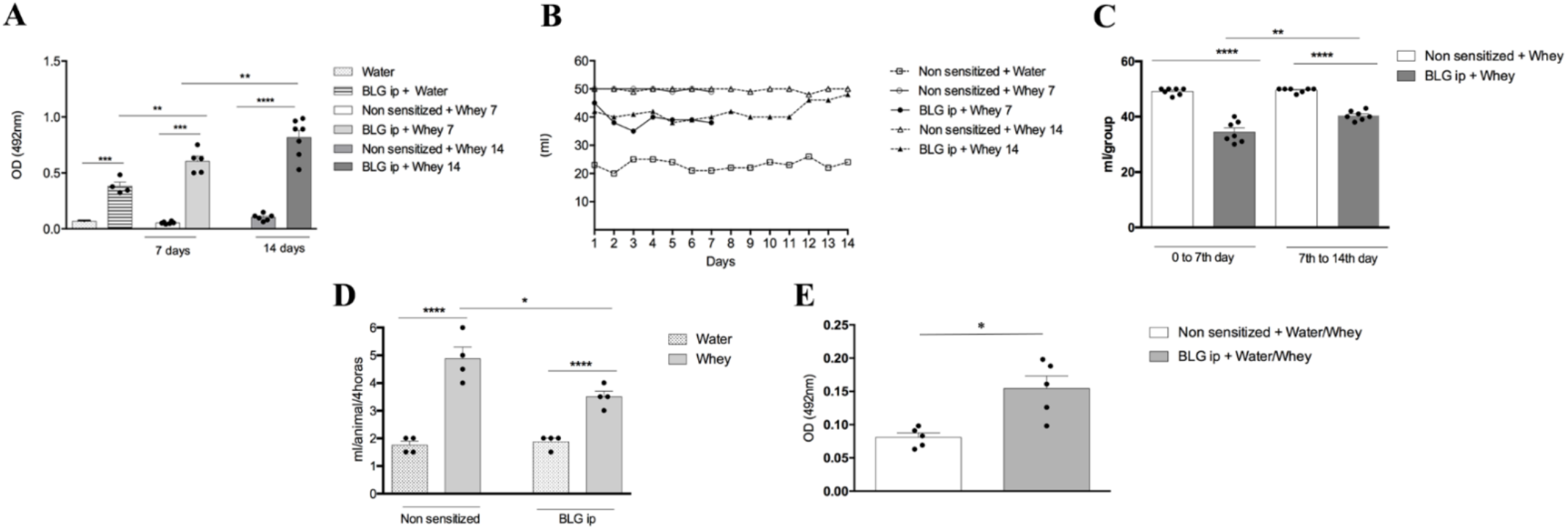
Beta-lactoglobulin induces partial aversion to consumption of whey-containing solution. A) anti-β-lactoglobulin IgE levels were measured in mice serum after oral challenge. B) Liquid intake among groups during oral challenge was measured daily. C) Liquid intake per group during two-time points: between day 0 until 7th and 7th until 14th day. D) Two-bottle preference test of control and sensitized BALB/c mice using Water or 20% Whey solution. Mice were sensitized twice with BLG/alum, on days 0 and 14. Control animals received only PBS. On day 21, animals were submitted to the preference test during 24h. E) Levels of IgE anti BLG were measured in sera after preference test. Data represent mean ± SEM.

To confirm the preference for the solution containing whey protein, we performed an experiment in a “two-bottle test" format (Mirotti et al., 2010), in which mice were offered the choice between ingesting water or a solution containing the allergen for which they were previously sensitized. The bottles were identical and there was a change of position between them to avoid preference guided by location. Interestingly, we found that sensitized mice preferred whey protein (Fig. 2B) even though they had high levels of anti-OVA IgE in the serum (Fig. 2C). Thus, the aversive behavior observed in BLG-allergic mice can be considered as partial since allergic mice still consumed more whey protein than control mice exposed to water and this effect was not definitive as previously shown for OVA-allergic animals (Cara et al., 1994).

### BLG possesses an anxiolytic effect

Anxiety is usually associated with the aversive behavior in mice allergic to OVA (Basso et al., 2003; Costa-Pinto et al., 2005). To investigate the association between aversion and anxiety in mice allergic to BLG, we performed a Zero Maze test in which each mouse is placed individually in the central area of the maze, where they can explore its arms freely for 15 minutes. The percentage of time spent in the open arms of the maze as well as the number of entries in the open arm are inversely proportional to the level of anxiety in the test. We found no difference among groups in the frequency of entries in the closed arms of the maze (Fig. 3A). However, non-sensitized mice that only ingested whey protein solution for 7 days showed more entries in the open arms of the maze, though no difference in time spent in the open arms among groups was observed (Fig3. B-C). Moreover, there was no difference in the distance traveled in the maze among groups (Fig3. D). To correlate behavior with activation of distinct brain areas, we measured c-Fos expression in the paraventricular neurons (PVN), an area related to anxiety, and found that non-sensitized mice drinking whey protein solution had lower levels of c-Fos expression in the PVN, revealing that fewer neurons were activated in this region (Fig 3E-F). Thus, it is likely that whey protein solution has an anxiolytic effect that is overcome by sensitization, since allergic mice had higher levels of c-Fos expression when compared to non-sensitized animals, but significantly lower levels than allergic mice that received water during the oral challenge period.

**Figure 3.**
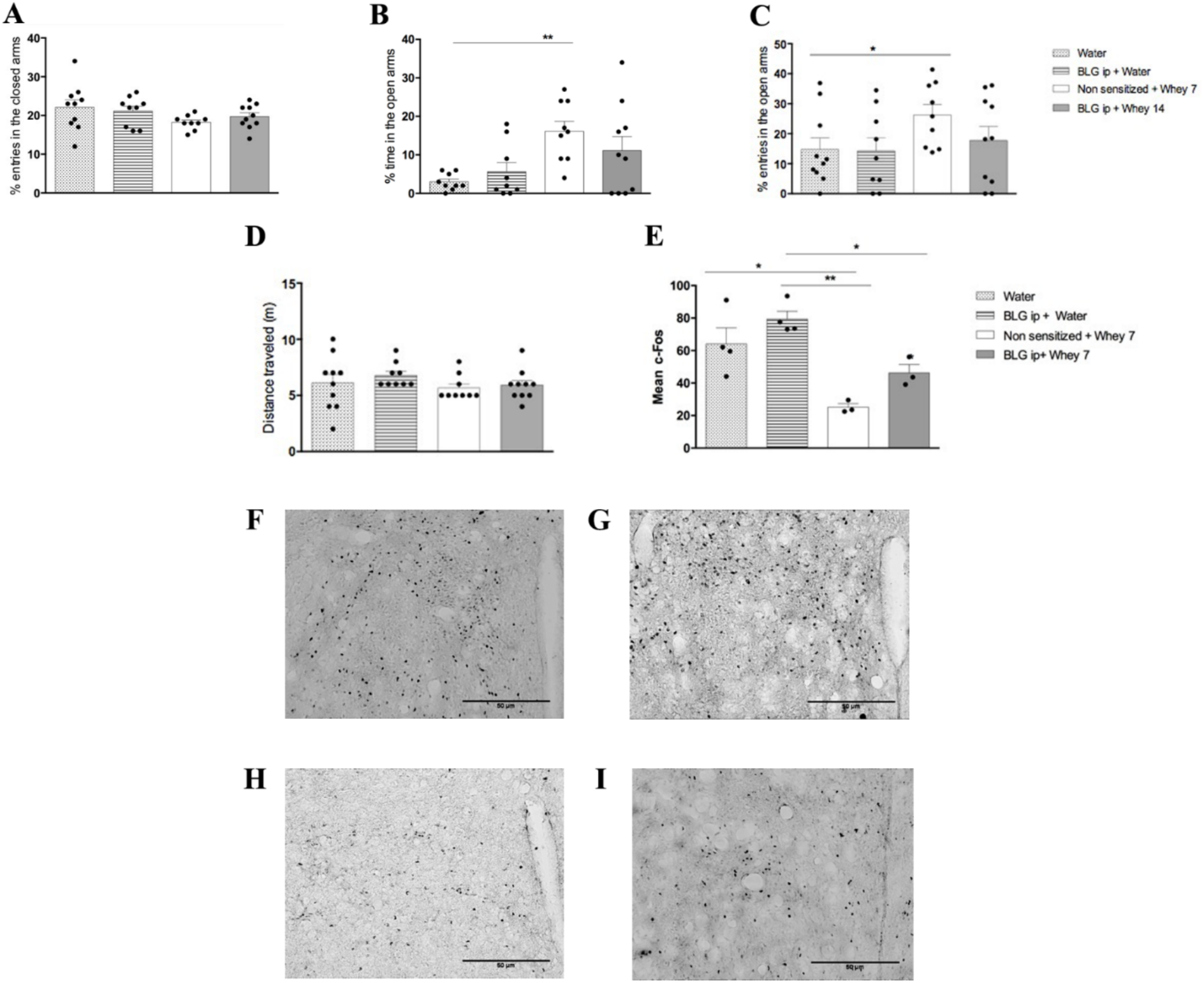
Analysis of the behavior of sensitized and non-sensitized mice in a Zero Maze test. BALB/c mice sensitized with BLG or non-sensitized, were placed one by one in a Zero Maze type labyrinth for 15 minutes. (A) the number of entries in the labyrinth's closed arms, (B) the percentage of time spent by the animals inside the open arms (C) the number of entries in the maze’s open arms (D) the total distance walked by the animals. Activated neurons in the paraventricular nucleus of the hypothalamus of BLG-sensitized mice. (E) Sensitized mice challenged or not with whey protein for 7 days pass through the Zero Maze labyrinth for 15 minutes at the end of the oral challenge. Ninety minutes after the labyrinth test, the animals were euthanized and perfused with saline and paraformaldehyde 4% and had their brains removed for subsequent slide preparation and immunohistochemical procedure. Neurons labeled with cFos in the PVN region were counted. * p <0.05, experiment with 6 animals per group. (F) Representative image of PVN area in non-sensitized mouse. (G) Representative image of PVN area in mice sensitized with BLG and treated with water in the oral challenge period. (H) cFos-tagged neurons in non-sensitized mice receiving whey protein solution, containing BLG, for 7 days. (I) cFos-marked neurons in mice sensitized and challenged with whey protein, for 7 days.

It is well known that mice have a strong preference for sweetened solutions (Cara et al., 1994, 1997). Thus, we hypothesized that BLG could produce the same effect. To investigate this, we performed a preference experiment in naive mice to determine whether mice preferred BLG over water, regardless sensitization. To ensure that our test was accurate, we chose to perform the preference test for 48 hours, measuring the consumption every 6 hours. We found that consumption of the BLG-containing solution was comparable with saccharin-containing solution, and significantly greater than water or OVA-containing solution (Fig. 4A). To ensure that purified BLG could reproduce the model previously performed with whey protein, we measure levels of anti-BLG IgE in mice sensitized to BLG and challenged with BLG rather than whey protein for 7 days and found that allergic mice produced higher levels of IgE than control mice (Fig. 4B). Moreover, we measured consumption of BLG-containing solution in these mice and found that aversion was present (Fig. 4C), indicating that mice challenged with BLG reproduce our findings with whey protein.

**Figure 4.**
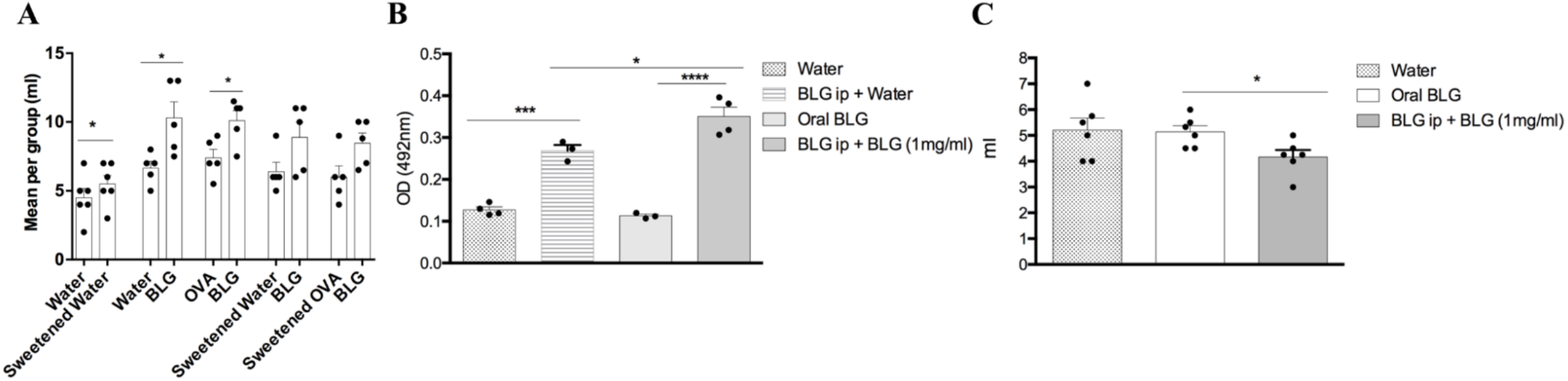
Preference test for solutions containing BLG. (A) Naive mice were kept individually separated and receiving the same amount of liquid so they could have access to two different sources of consumption. Mice were divided into following groups: water x BLG (1mg / ml), OVA x BLG (1mg / ml), Water sweetened (1% saccharin sodium) x BLG (1mg / ml) and OVA sweetened / ml). The graph shows the consumption of preference test. * p <0.05. Bars represent mean + e.p.m. Experiment made with 5 mice per group. (B) Levels of total IgE in serum samples after oral challenge (C) Liquid intake among groups during oral challenge was measured daily.

Taken together, the less activation of brain areas related to anxiety and the strong preference to both whey protein and BLG over water or OVA-containing solution, suggest that BLG may activate reward-associated brain areas that potentially overcome the aversive behavior.

### BLG activates the brain reward system

Since naive mice had a preference for BLG-containing solution, we used a classic OVA allergy model, in which aversion is clearly established (Cara et al., 1994; Saldanha et al., 2004), to verify whether the presence of BLG altered the aversive behavior to OVA. As expected, we found that mice sensitized to and challenged with OVA alone or both OVA and BLG had high levels of anti-OVA IgE (Fig. 5A). Furthermore, mice sensitized to OVA and challenged with OVA consumed less OVA, which characterizes an aversive behavior, but mice challenged with a solution containing BLG alone or both OVA and BLG, consumed significantly more solution than mice only exposed to OVA (Fig. 5B), suggesting that BLG triggered an effect similar to that observed with sweetened solutions, which may indicate an activation of brain-related reward areas.

**Figure 5.**
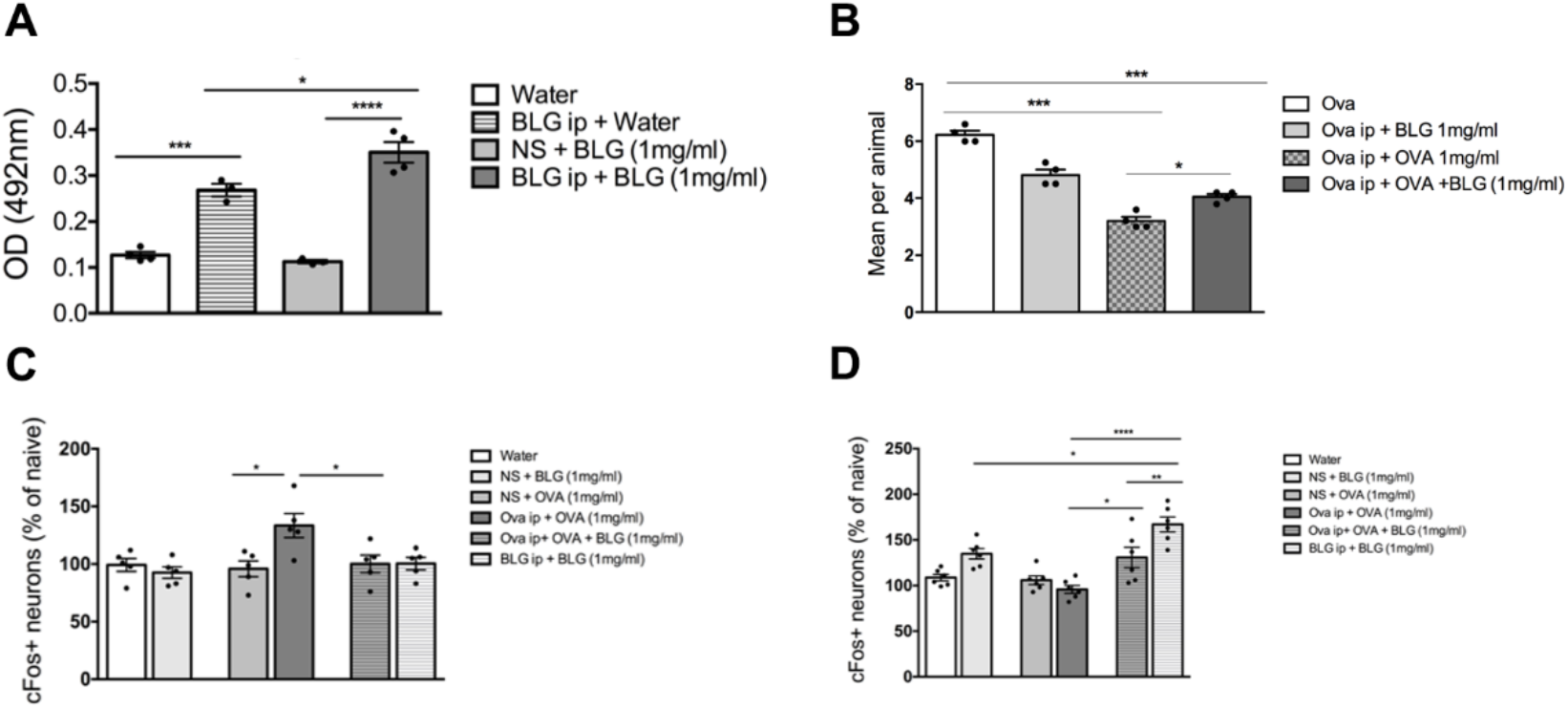
BLG modifies aversion to OVA in a food allergy model to OVA. BALB/c mice were sensitized i.p with OVA and challenged with OVA (1mg / ml) for three days or with solution containing OVA and BLG (1mg/ml). The group NS + OVA was not sensitized and received only oral OVA, as experimental control. (A) OVA-specific IgE levels expressed as absorbance at 492nm. (B) Variation of body weight per group during oral challenge (C) Average of liquid consumption per animal. Results are expressed as mean + e.p.m. p <0.05. (D) Evaluation of cFos expression in mice. BALB/C mice were divided into naive (non-sensitized) mice that ingested OVA (NS + OVA); non-sensitized mice that ingested BLG (NS + BLG); mice that were sensitized i.p for OVA and challenged orally with OVA (OVA ip + OVA) and a group that were sensitized to OVA and challenged with a mixture of BLG and OVA (1mg/ml) * p <0.05.

To investigate the brain areas that could be associated with this behavior, we evaluated two brain areas based on c-Fos expression: the central nucleus of the amygdala (CeA), related to anxiety and protective behaviors, and the nucleus accumbens (NAc), responsible for a positive reward system. We found that mice sensitized to and challenged with OVA showed more activation of neurons from the CeA. No activation was observed in mice sensitized to BLG and challenged with BLG. Surprisingly, mice that were sensitized to OVA and challenged with a solution containing BLG alone or both OVA and BLG showed less activated neurons in this area (Fig. 5C). On the other hand, BLG consumption induced a strong activation of neurons from the NAc, even when mice were challenged with OVA, indicating that BLG indeed activates the reward system, which is likely the reason why mice allergic to OVA lose the aversive behavior when exposed to BLG (Fig. 5D).

## DISCUSSION

β-lactoglobulin (BLG) is considered to be the most immunogenic protein among cow’s milk allergens (Lindholm Bøgh et al., 2013). Among the many clinical alterations typical of food allergy, behavioral changes such as aversion and anxiety have been described in allergic individuals (Costa-Pinto and Basso, 2012). In mice, the connection between food allergy and behavior has been shown to be allergen specific because mice sensitized to peanut or wheat when offered with a mixture of the grains *in natura* chose to ingest grains that there were not previously exposed (Teixeira G. 1995). The same behavioral changes were also demonstrated by using OVA as the allergenic protein (Basso et al., 2003; Cara et al., 1994). However, the mechanisms underlying these behavioral alterations in food allergy are still elusive. Thus, in the present study, we sought to investigate whether behavioral alterations occur in a model of food allergy induced by BLG, and the neuro-immunological mechanisms involved in this effect.

We showed that BALB/c mice sensitized to BLG and later challenged with a 20% whey protein (80% BLG) solution developed a food allergy characterized by weight loss and small intestine mucosa inflammation, which was associated with increased frequency of eosinophils, goblet cells and intraepithelial lymphocytes. In addition, allergic mice had higher secretory IgA levels after being challenged with whey protein-containing solution. As for the cytokine profile, IL-4 and IL-5 were increased in allergic mice accompanied by a decrease in IL-10. Moreover, BLG allergic mice had elevated levels of serum anti-BLG IgE, but contrary to previous reports on OVA-induced food allergy (Batista et al., 2014), prolonged exposure to BLG (14 days) further increased IgE levels as compared to 7 days of oral challenge. Importantly, while OVA allergic mice showed the characteristic aversive behavior when exposed to OVA, mice allergic to BLG had only a partial aversion to BLG, which was abrogated by day 12 post challenge despite the progressive increase of anti-BLG IgE. This is intriguing because it has been described that the aversive behavior depends on the production of IgE and that the depletion of IgE by anti-IgE antibodies prior to oral challenge prevents the development of aversion (Basso et al., 2003). Moreover, this effect was dependent on mast cell degranulation, because cromolyn, a mast cell stabilizer, completely blocked c-Fos expression in brain areas associated with anxiety and aversion, such as the paraventricular nucleus of the hypothalamus (PVN) and in the central nucleus of the amygdala (CeA), and mice treated with cromolyn lost the aversive behavior to OVA (Costa-Pinto et al., 2007). Thus, BLG may activate the reward system, which overcomes the aversive behavior induced by allergy. We confirmed this hypothesis by showing that (i) consumption of the BLG-containing solution was comparable with a sweetener-containing solution, and significantly greater than water or OVA. Taste preference for sweet solutions are associated with the activation of the reward system (Low et al., 2014); (ii) unlike OVA, BLG decreased c-Fos expression in neurons from the PVN and CeA, indicating that BLG has anxiolytic properties; and (iii) BLG increased c-Fos expression in neurons from the nucleus accumbens (NAc), a critical area belonging to the reward system that is involved in processing incentive salience, pleasure, positive reinforcement and reinforcement learning (Goto and Grace, 2005). Thus, different allergens induce different taste preferences and the outcome is related to neuronal components triggered by the allergens.

Although we have clearly shown that BLG, unlike OVA, activates neurons from reward brain areas, the mechanism underlying this effect remains unknown. Enzymatic digestion of natural proteins, including those derived from milk, generates a range of bioactive peptides that may interact with the nervous system and exhibit anxiolytic activity. Alpha-lactotensin (His-Ile-Arg-Leu, HIRL) for example, is a peptide that was isolated from chymotrypsin digestion of BLG (Yamauchi et al., 2003) and has anxiolytic effect in mice in behavioral tests using a high cross maze (Hou et al., 2011). It is interesting that some bioactive peptides derived from food components such as whey proteins have inhibitory activity for angiotensin converting enzyme (ACE) that regulates blood pressure. Inhibition of this enzyme drives anxiolytic effects in rats subjected to behavioral tests (Welderufael et al., 2012). Thus, it is possible that BLG acts as an anxiolytic through the action of its peptides.

Another plausible possibility is the communication between the digestive system and the brain. It is known that the aversive behavior to OVA, as mentioned above, is associated with neuronal activation in the PVN and CeA, which are brain areas related to emotional and affective behavior. Moreover, PVN and CeA are among the main regions containing neurons expressing the corticotropin-releasing hormone (CRH), which is a key peptide in controlling behavioral, neuroendocrine and autonomic responses to stress, anxiety and depression (Arborelius et al., 1999; Heinrichs et al., 1995; Holsboer and Ising, 2008). Consistent with this, mice with food allergy display higher levels of anxiety and increased serum corticosterone levels. Thus, the CRH-corticosterone axis may play an important role in the anxiety observed during food aversion. Moreover, stabilization of mast cells by cromolyn prevented the aversive behavior to OVA (Costa-Pinto et al., 2007), suggesting that the early phase of an immediate allergic response is critical for the aversion development. Importantly, mast cells secrete a variety of mediators including growth factors, cytokines, histamine and serotonin (Galli et al., 2008; Kalesnikoff and Galli, 2008). Interestingly, mast cells are closely apposed to nerve endings from the vagal nerve in both humans and rodents (Stead et al., 1989) giving anatomical support for mast cell-secreted mediators to interact with neurons. Consistent with this, treatment of neonatal mice with capsaicin, a neurotoxin derived from chilli pepper that selectively promotes the dysfunction of the sensory fibers named C-fibers, blocked c-Fos expression in the PVN and reduced food aversion to OVA-sensitized mice (Basso et al., 2004, 2001). Furthermore, selective antagonism of the serotonin 5-HT3 receptors, which are expressed in C-fibers (Lang et al., 2006) decreased the aversive behavior in rats (Zarzana et al., 2009). Thus, food aversion may occur due to the release of mast cell factors upon IgE/allergen binding that stimulate sensory fibers of the vagal nerve, which in turn drive these peripheral informations to the central nervous system. Additionally, since IgE receptor (FcεRI) has been detected on sensory neurons from mice (Andoh and Kuraishi, 2004; van der Kleij et al., 2010), a direct binding of IgE on vagal nerve sensory fibers may represent an alternative neuronal activation independent of mast cells. However, the circuits involved in gut-activated brain reward system have remained elusive until recently in which Han et al demonstrated that optical activation of gut-innervating vagal sensory neurons recapitulated the classical effects of stimulating brain reward neurons (Han et al., 2018). Thus, we believe that the reward behavior induced by BLG is a consequence of the stimulation of specialized sensory fibers in the vagal nerve responsible for activating brain areas associated with pleasure and reward such as the NAc.

In summary, we showed that the aversive behavior observed in food allergy to OVA does not apply to all antigens. Mice allergic to BLG, although presenting high serum IgE levels and several signs of inflammation in the intestinal mucosa, ingested solutions containing the allergen even when they were offered other options. Our data suggest that BLG has special characteristics that could be related to its taste or to neurologically active peptides present in its structure, that can subvert the evolutionarily preserved behavior of rejection to potentially toxic substances triggered by allergic reactions dependent on IgE.

## Author Contributions

L.L. and H.C.A. performed and followed all the experiments, also wrote the paper. D.S.R., J.L.A., M.C.C., B.K.S.M. and T.G.M. contributed in the analysis of the experiments and elaboration of graphs, M.A.M. and L.M.P. performed cFos staining and analyses, supervised by B.R.S; L. A. B and J.A.S.G contributed in the behavior tests, supervised by D. C. A.; D.C.C. contributed to the histological analysis. A.C.G.S. and A.M.C.F. contributed in the delineation of the experiments and supervision of the work.

## Data availability statement

All datasets generated or analyzed during the current experimental study are available from the corresponding author on reasonable request.

## Ethical approval and informed consent

This project was approved by the Ethics Committee on Animal Use at Federal University of Minas Gerais (CEUA/UFMG) with protocol no. 114/19, related to the present study is in agreement with the Ethical Principles in Animal Experimentation.

## Funding

This work was supported by Conselho Nacional de Desenvolvimento Científico e Tecnológico (CNPq), Fundação de Amparo à Pesquisa do Estado de Minas Gerais (FAPEMIG), and Coordenação de Aperfeiçoamento de Pessoal de Nível Superior (CAPES).

## Conflict of Interest Statement

The authors declare that the research was conducted in the absence of any commercial or financial relationships that could be construed as a potential conflict of interest.

## Acknowledgements

The authors are thankful to Ilda Marçal de Sousa and Hermes for their excellent work taking care of our animal facility. We thank Dr. Rafael Rezende (Brigham and Women’s Hospital, Harvard Medical School) for discussing and providing inputs to the manuscript). This study was supported by a Grant from Conselho Nacional de Desenvolvimento Científico e Tecnológico (CNPq), Brazil. Some of the authors are recipients of scholarships (L Lemos; MCC Canesso) and fellowships (A.M.C.Faria, D.C.Cara) from Conselho Nacional de Desenvolvimento Científico e Tecnológico (CNPq), Brazil.

## Notes

### Competing Interest Statement

The authors have declared no competing interest.

